# Chromatin features define adaptive genomic regions in a fungal plant pathogen

**DOI:** 10.1101/2020.01.27.921486

**Authors:** David E. Cook, Martin Kramer, Michael F. Seidl, Bart PHJ Thomma

**Author notes:** Corresponding authors: David E. Cook,; Bart PHJ Thomma.

## Abstract

Understanding the complex information stored in a genome remains challenging since multiple connected regulatory mechanisms act at various scales to determine function. Increased comprehension of genome function at scales beyond contiguous nucleotides will help understand genetic diseases, the emergence of pathogenesis, and more broadly the genomics of adaptation. Here we report the analysis of DNA methylation, histone modification, and DNA accessibility in the plant pathogenic vascular wilt fungus *Verticillium dahliae*. Functional analysis details that DNA methylation is restricted to repetitive elements, such as transposable element DNA, but interestingly only some repetitive DNA is methylated. This incomplete DNA methylation is associated with repetitive DNA residing in specific compartments of the genome that were previously defined as Lineage-Specific (LS) regions. These regions are hypervariable between *V. dahliae* isolates and contain genes that support host colonization and adaptive traits. LS regions are associated with H3 Lys-27 methylated histones (H3K27me3), and repetitive DNA within LS regions are more transcriptionally active and have increased DNA accessibility, representing a hybrid chromatin state when compared to repetitive regions within the core genome. We used machine learning algorithms trained on epigenetic and DNA accessibility data to predict LS regions with high recall, identifying approximately twice as much LS DNA in the *V. dahliae* genome as previously recognized. Collectively, these results characterize LS regions in an intermediate chromatin state and provide evidence that links chromatin features and genome architecture to adaptive regions within the genome.

## INTRODUCTION

Genomes are not randomly organized and comprise complex information beyond their linear nucleic acid sequence ^1^. While scientific understanding of genome biology continues to grow, significant efforts in the past decade have focused on sequencing new species and additional genotypes of those species ^2^. However, there is a great need to decode the complex information stored in these genomes, to understand genomic responses over various time scales, and ultimately to more fully understand how genotypes lead to phenotypes. With the growing number of high-quality, highly contiguous genome assemblies it is possible to analyze genome organization into chromosomes at high resolution ^3^. Present day genome organization reflects evolutionary solutions to the challenges of information processing and adaptation; a genome must faithfully pass vast amounts of information across cell-cycles and reproduction, packaged into limited physical space, while achieving correct access to the information in response to developmental, environmental or chemical signals. In addition, there needs to be appreciable stochastic genetic variation to ensure that phenotypic variation is present for unknown future events. Organisms undergoing mainly asexual reproduction face an additional evolutionary constraint as they must generate this genetic variation in the absence of meiotic recombination ^4^. Many economically important fungal plant pathogens are either asexual or undergo more frequent asexual reproduction compared to sexual reproduction ^5^. Interestingly, fungal pathogens are subject to additional evolutionary pressure from their hosts, as host-pathogen interactions create dynamical systems with shifting, yet near-constant selective pressure on the two genomes ^6^. These attributes make plant-fungal interactions a particularly interesting system to study aspects of genome evolution and genome organization ^7,8^.

Plant invading microbes use effectors to suppress, avoid or mitigate the plant immune system ^9,10^. Plants in-turn use a variety of plasma-membrane bound and cytoplasmic receptors to recognize invasion, through recognition of the effector or its biochemical activity, creating a strong selective pressure on the microbe to modify the effector or its function to alleviate recognition ^11,12^. The plant pathogenic fungus *Verticillium dahliae* causes vascular wilt diseases on hundreds of plant hosts. *V. dahliae* is presumed asexual and generates genomic diversity in the absence of sexual recombination through large-scale chromosome re-arrangements and segmental duplications ^13–16^. The regions undergoing such duplications and re-arrangements are hypervariable between *V. dahliae* isolates, and consequently have been referred to as Lineage-Specific (LS) regions. These LS regions are enriched for *in planta* expressed genes and harbor many effector genes contributing to host infection ^14,17,18^. Similar non-random genomic arrangement of effectors have been reported across diverse plant pathogenic fungal and oomycete genomes ^14,19–25^. One summary of these observations is referred to as the two-speed genome, in which repeat-rich regions harboring effectors evolve more rapidly than genes outside these regions ^26^.

Previous research in various plant-associated fungi has established a link between posttranslational histone modifications and transcriptional regulation of adaptive trait genes. These genes include effectors that facilitate host infection, and secondary metabolite (SM) clusters that code for genes that produce chemicals important for niche fitness ^27^. By removing or reducing enzymes responsible for particular repressive histone modifications, such as di- and trimethylation of Lys9 and Lys27 residues of histone H3 (H3K9me2/3 and H3K27me2/3), a disproportionally high number of effector and SM cluster genes are derepressed, although a direct role of these marks in transcriptional control was not demonstrated ^28–30^. However, evidence from the fungus *Epichloe festucae* that forms a mutualistic interaction with its grass host *Lolium perenne* indicates that direct transcriptional regulation through histone modification dynamics is possible ^31^. Although there are clear indications that the epigenome (i.e. heritable chemical modifications to DNA and histones not affecting the genetic sequence) plays a role in adaptive gene regulation, additional evidence is needed to fully understand this phenomenon.

Epigenetic modifications influence chromatin structure, defined as the DNA-RNA-protein interactions giving DNA physical structure in the nucleus ^32,33^. This physical structure affects how DNA is organized in the nucleus and DNA accessibility. Methylation of H3K9 and H3K27 are hallmarks of heterochromatin; DNA that is tightly compacted in the nucleus ^34–37^. H3K9 methylation is not only associated with controlling constitutive heterochromatin, but also tightly linked with DNA cytosine methylation (mC), which serves as an epigenetic mark contributing to transcriptional silencing ^38^. A single DNA methyltransferase gene, termed *Dim2*, performs cytosine DNA methylation in the saprophytic fungus *Neurospora crassa* ^39^. Histone methylation at H3K9 directs DNA methylation by DIM2 through another protein, termed heterochromatin protein 1 (HP1), which physically associates with both DIM2 and H3K9me3 ^40,41^. Some fungi possess a unique pathway to limit the expansion of repetitive DNA such as transposable elements through repeat-induced point mutation (RIP), a mechanism that specifically mutates repetitive DNA in the genome during meiosis and induces heterochromatin formation ^42,43^. The mutations occur at methylated cytosines resulting in conversion to thymines (C to T mutation)^44^. H3K27 methylation is associated with heterochromatin that is thought to be more flexible in its chromatin status and exist as bivalent chromatin that may be either transcriptionally repressed or active depending on developmental stage or environmental cues ^45–48^. The deposition of H3K27me3 is controlled by a histone methyltransferase that is a member of a complex of proteins termed Polycomb Repressive Complex 2 (PRC2), with orthologs of the core machinery present across many eukaryotes ^36,49^.

In addition to heterochromatin playing a role in transcriptional regulation in filamentous fungi, epigenetic marks contributing to chromatin may influence genome evolution ^50^. In *N. crassa*, DNA is physically arranged in the nucleus corresponding to heterochromatic and euchromatic domains, with strong inter- and intra-heterochromatin DNA-DNA interactions reported ^51,52^. Recent experimental evidence using *Zymoseptoria tritici*, a fungal pathogen of wheat, suggests that H3K27me3 promotes genomic instability ^53^. In the oomycete plant pathogens *Phytophthora infestans* and *Phytophthora sojae* a clear association exists between gene-sparse and transposon-rich regions of the genome and the occurrence of adenine N6-methylation (6mA) ^54^. Collectively these examples point towards an unexplained connection between the epigenome, genome architecture, and adaptive evolution. To examine the hypothesis that epigenetic modifications influence the adaptive LS regions of *V. dahliae*, we performed a series of genetic, genomic, and machine learning analyses to characterize these regions in greater detail.

## RESULTS

### DNA cytosine methylation occurs at transposable elements

To understand the role of DNA methylation in *V. dahliae*, whole-genome bisulfite sequencing, in which unmethylated cytosine bases are converted to uracil while methylated cytosines remain unchanged ^55,56^, was performed in the wild-type and a heterochromatin protein 1 deletion mutant (*Δhp1*). The overall level of DNA methylation in *V. dahliae* is low, with an average weighted methylation percentage (calculated as the number of reads supporting methylation over the number of cytosines sequenced) at CG dinucleotides of 0.4% (Table 1). The fractional CG methylation level (calculated as the number of cytosine positions with a methylated read over all cytosine positions) is higher, averaged to 9.7% over 10 kb windows. Weighted and fractional cytosine methylation (mC) levels are statistically significantly higher in the WT compared to the *Δhp1* mutant for all cytosine contexts (Table 1, Supplemental Fig. S1A and B). This result is consistent with the requirement of HP1 for DNA methylation in *N. crassa* ^40^. To understand DNA methylation in the context of the functional genome, DNA methylation was analyzed over genes, promoters, and transposable elements (TE). Despite statistically significant differences between WT and *Δhp1* for gene and promoter methylation, the bisulfite sequencing data shows virtually no DNA methylation at these two features (Fig. 1A). We attribute the difference to a marginal set of elements having a real difference between the genotypes, but the biological significance is likely negligible (Fig. 1A). In contrast, there is a much higher degree of methylation, and a notable difference between wild-type and *Δhp1* methylation levels at TEs (Fig. 1A, bottom panel), with the average CG methylation level being five times higher in the wild-type strain.

**Figure 1.**
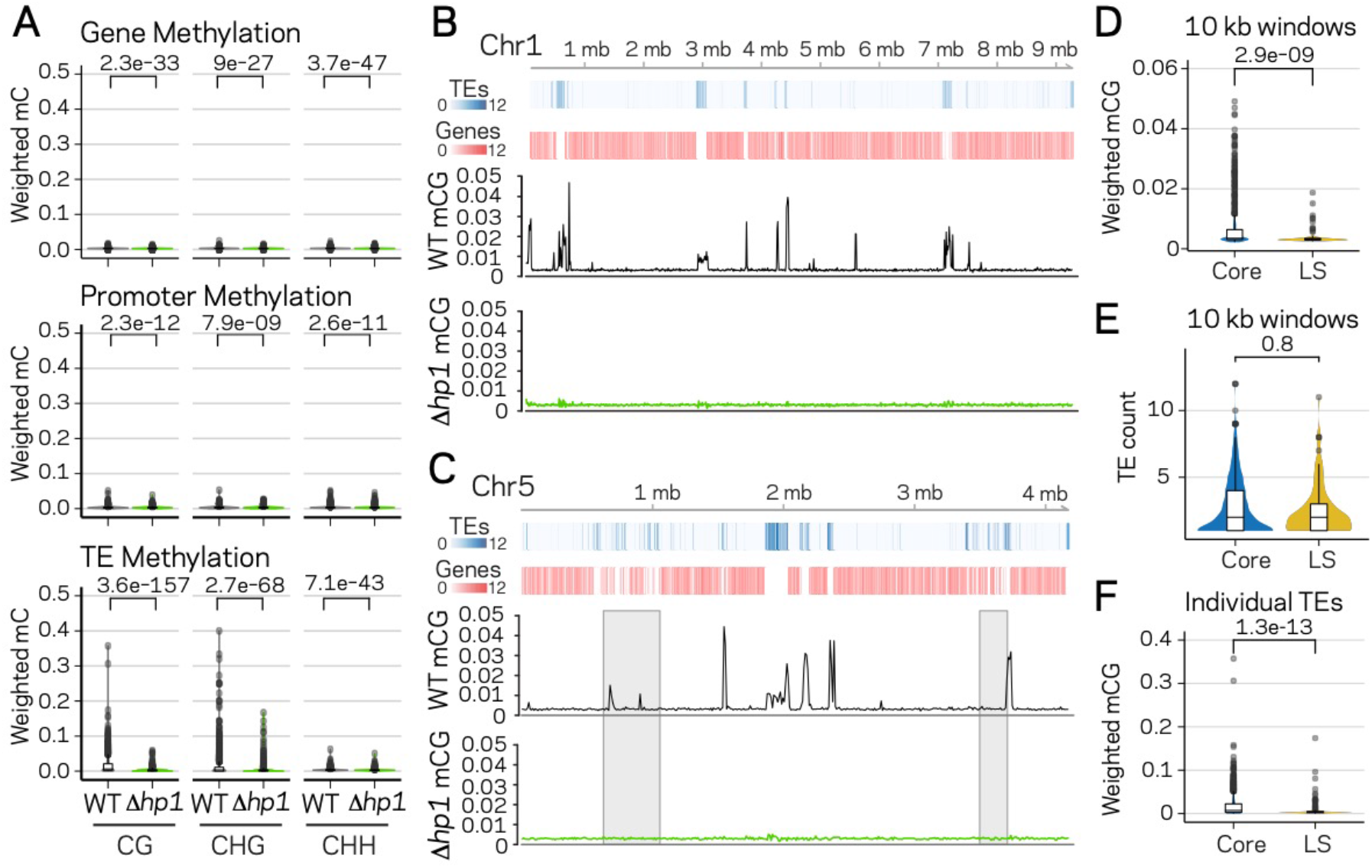
DNA methylation is only present at transposable elements, but not at those present in LS regions. (A) Violin plot of the distribution of DNA methylation levels quantified as weighted methylation over Genes, Promoters and TEs. Cytosine methylation was analyzed in the CG, CHG and CHH sequence context. Methylation was measured in the wild-type (WT) and heterochromatin protein 1 knockout strain (Δ*hp1*). (B, C) Whole chromosome plots showing TE and Gene counts (blue and red heatmaps) and wild-type (black lines) and Δ*hp1* (green line) CG methylation as measured with bisulfite sequencing. Data is computed in 10 kilobase non-overlapping windows. (C) Two previously defined LS regions (Faino *et al.* 2016) are highlighted by grey windows. (D) Violin plot of weighted cytosine methylation in 10 kb windows broken into core versus LS location (E) Same as D but plots are for the counts of TEs per 10 kb window. (F) Same as in D but methylation levels were computed at individual TE elements. Statistical differences for indicated comparisons were carried out using non-parametric Mann-Whitney test with associated p-values shown.

**Table 1.** Summary of DNA methylation in *Verticillium dahliae* wild-type (WT) and heterochromatin protein 1 deletion mutant (*Δhp1*) as measured by whole genome bisulfite sequencing calculated over 10 kb non-overlapping windows.

| Genotype | Avg. Weighted mCG | Avg. Weighted mCHG | Avg. Weighted mCHH | Avg. Fraction mCG | Avg. Fraction mCHG | Avg. Fraction mCHH |
| --- | --- | --- | --- | --- | --- | --- |
| WT | 0.0040 | 0.0037 | 0.0034 | 0.097 | 0.097 | 0.088 |
| $\Delta hp1$ | 0.0030 | 0.0030 | 0.0032 | 0.082 | 0.083 | 0.079 |

Avg. Weighted, The average of total methylated cytosines in a given context divided by total cytosines in that context in a 10 kb windows; Avg. Fraction, The total cytosines positions with a read supporting methylation divided by total cytosines in a specific context in a 10 kb window; mCG, methylated cytosine residing next to a guanine; mCHG, methylated cytosine residing next to any base that is not a guanine next to a guanine; mCHH, methylated cytosine residing next to any two bases that are not a guanines.

To further analyze DNA methylation levels and confirm that the low DNA methylation levels in the wild-type strain are indeed different than those in *Δhp1,* CG DNA methylation levels were plotted in 10 kb windows across individual chromosomes. These plots clearly show that DNA methylation is not continuously present across the *V. dahliae* genome, and DNA methylation is significantly different between wild-type and *Δhp1* (Fig. 1B, C). Furthermore, regions in the genome with higher densities of TEs and lower gene numbers have higher levels of DNA methylation, consistent with the global DNA methylation summary (Fig. 1B and C). Interestingly, these results show that while DNA methylation is only present at TEs, not all TEs are methylated, a phenomenon that was previously described as ‘non-exhaustive’ DNA methylation ^57^. To further understand this phenomenon, we sought to identify discriminating genomic features that could account for some TEs not being methylated. The whole-chromosome methylation data suggested a lack of DNA methylation at previously identified LS regions (Fig. 1C, grey windows). These LS regions were previously detailed for *V. dahliae,* and are characterized as regions that are highly variable between isolates of the species, are enriched for actively transcribed TEs, and contain an increased proportion of genes involved in host virulence ^13–15^. Thus, we tested if DNA sequences at LS regions are less frequently methylated by comparing weighted mCG levels in 10 kb bins containing at least one TE for core versus LS regions. This analysis showed significantly more DNA methylation for core bins, which cannot be accounted for by a simple difference in the number of TEs in the core and LS regions analyzed (Fig. 1D and E). Higher CG methylation levels also hold true when analyzed at the level of individual TE elements (Fig. 1F, numbers of elements in Supplemental Table S1). Collectively, these analyses demonstrate that DNA methylation occurs almost exclusively at TEs and, importantly, that not all TEs are methylated. This observation can in part be explained by mCG differences for TEs in the core versus LS regions.

### Transposable element classes have distinct profiles for genomic and epigenomic features

To understand the functional status of the various TEs in the genome, DNA-histone modification location data were collected using chromatin immunoprecipitation followed by sequencing (ChIP-seq) against H3K9me3 and H3K27me3, which allows for the identification of DNA interacting with these modified histones. Characteristics of TE sequence, such as GC percentage, composite RIP index (CRI), and TE age, estimated as the Jukes-Cantor distance to the consensus sequence of the specific TE family, were calculated (see methods). To further classify genomic regions as eu- or heterochromatic, we performed an assay for transposase accessible chromatin and sequencing (ATAC-seq) ^58^. This method uses a TN5 transposase to restrict physically accessible DNA in the nucleus and tags the DNA ends with oligonucleotides for downstream sequencing. Transcriptional activity was assayed using RNA-sequencing. To analyze all of these TE characteristics (variables) at once, dimensional reduction with principle component analysis (PCA) was employed, which facilitates data interpretation on two-dimensions to identify important variables and their relationships within large datasets. The individual TEs were grouped into four broad classes (Type I DNA elements and Type II LTR, LINEs, and Unspecified elements) and analyzed for each measured variable. The first dimension of PCA shows the largest separation of the data points and variables, and largely separates the data based on euchromatin versus heterochromatin features (Fig. 2A, PC1). This is seen by the variables ATAC-seq, %GC, RNA-sequencing, H3K9me3 ChIP, CRI and DNA methylation (mCG) being furthest separated along the x-axis (Fig. 2A). Open chromatin features such as higher ATAC-seq, %GC, and transcriptional activity are positive on the x-axis, with small angles between the vectors, indicating correlation among those variables. Conversely, features associated with heterochromatin, such as H3K9me3 association, DNA methylation and indication of RIP (CRI) are all negative on the x-axis, and the position of their vectors indicates correlation among these variables, and negative correlation to the euchromatin features (Fig. 2A). The second axis discriminates elements based on their H3K27me3 profile and sequence characteristics such as Jukes Cantor (TE age), Identity and Length (Fig. 2A). For the individual element classification, there is a stronger association for the LTR and Unspecified elements with the heterochromatin features (Fig. 2A, grey and red ellipse extending along negative x-axis). Collectively, this multivariate description of TEs identifies those that are more transcribed and open as having lower association with H3K9me3, mCG, and RIP mutation. There are statistically significant differences between the TE types for each of these variables (Supplemental Table S2), and the LTR elements have the highest levels of H3K9me3 and mCG, along with the highest CRI values and lowest %GC, consistent with the mechanistic link between the four variables (Fig. 2B). Interestingly, a bimodal distribution occurs for %GC and CRI within the LTR and Unspecified elements, indicating that some of the LTR elements have undergone RIP and are heterochromatic, while other elements have not been subject to this mechanism (Fig. 2B). This delineation occurs for the Unspecified and LTR elements with a %GC sequence content less than approximately 40%, which have positive CRI values and high H3K9me3 signal (Fig. 2C). A similar distinction is seen with ATAC-seq data that show a clear break around 40% GC content, and elements below this have lower ATAC-seq signal and higher H3K9me3 signal (Fig. 2D). These trends are not observed for the LINE and DNA elements (Supplemental Fig. S2). These results suggest that LTR and Unspecified TE elements exist in two distinct chromatin states in the genome.

**Figure 2.**
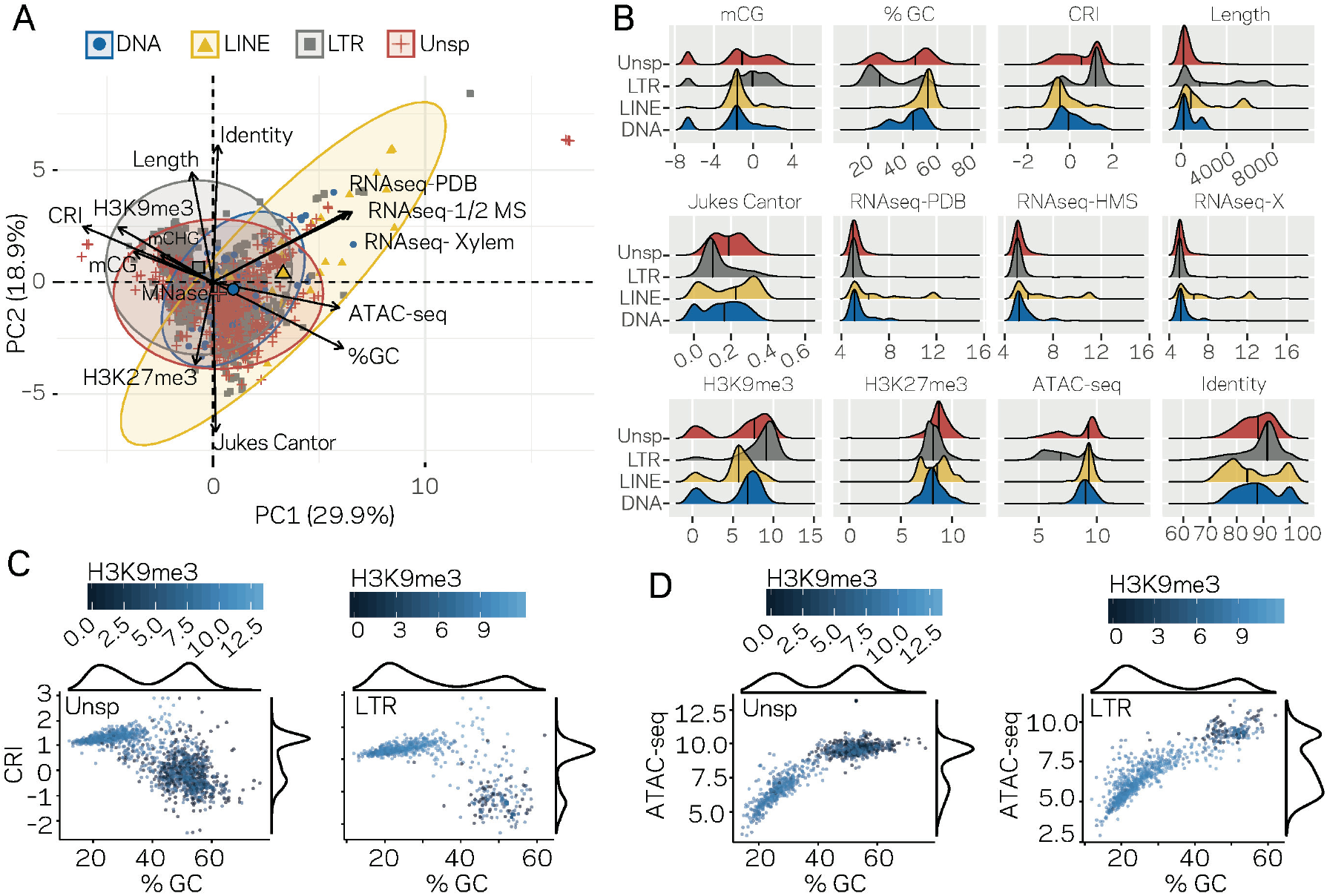
Individual TE families have distinct epigenetic and physical compaction profiles. (A) Principle component analysis for 14 variables measured for each individual TE. Each vector represents one variable, with the length signifying the importance of the variable in the dimension. The relationship between variables can be determined by the angle connecting two vectors. For angles <90^0^, the two variables are correlated, while those >90^0^ are negatively correlated. Each individual element is shown and highlighted by color and symbol as indicated by the key. Colored ellipses show the confidence interval for the four families along with a single large symbol to show the mean position for the four families. mCG, weighted CG DNA methylation; mCHG, weighted CHG DNA methylation; CRI, Composite RIP index; %GC, percent GC sequence content; Identity, Nucleotide identity as percent identity to the consensus TE sequence of a family; Length, element length; Jukes Cantor, Jukes Cantor corrected distance as proxy of TE age; RNAseq, RNA-sequencing reads from (PDB), half strength MS (HMS) or tomato xylem sap (Xylem) grown fungus expressed as variance stabilizing transformed log2 values (see methods for details); H3K9me3, log2 (TPM+1) values of mapped reads from H3K9me3 ChIP-seq; H3K27me3, log2 (TPM+1) values of mapped reads from H3K27me3 ChIP-seq; ATAC-seq, log2 (TPM+1) values for mapped reads from Assay for transposase accessible chromatin. (B) Ridge plots showing the distribution of the individual TE families per variable. The median value is shown as a solid black line in each ridge. Variables same as in A except for mCG, log2(weighted cytosine DNA methylation + 0.01). (C) Scatter plot for %GC versus CRI values for individual TE elements shown as points. The two plots are for TEs characterized as Unspecified (Unsp) or LTR, labeled in the upper left corner. Each point is colored according to log2 (TPM+1) values from H3K9me3 ChIP-seq, scale shown above each plot. A density plot is shown for both variables on the opposite side from the labeled axis. (D) Same as in C, but the y-axis is now showing the log2 (TPM+1) values from ATAC-seq.

### Transposable element location significantly influences the epigenetic and DNA accessibility profile

To further dissect the relationship between epigenetic modifications, chromatin status and genomic location, pair-wise comparisons were made for all TEs in core versus LS regions. All measured variables, except TE length, are significantly different for TEs in the core versus LS regions (Supplemental Fig. S3). Further division of the TEs indicated that the LTR and Unspecified elements showed the greatest differences for core versus LS measurements (Fig. 3A), demonstrating that the major driver of core versus LS differences are driven by the LTR and Unspecified elements. The bimodal distribution for %GC, CRI, H3K9me3, and ATAC-seq can be accounted for in part by core versus LS separation (Fig. 3B, red versus grey). Collectively, the status of the LS TE elements can be characterized as devoid of DNA and H3K9 methylation, low RIP signal, generally higher than 50% GC content, higher levels of H3K27me3, more open with ATAC-seq signal, and higher transcription levels (Fig. 3D). The core versus LS location is not sufficient to fully explain the chromatin status, as there are many elements located in the core genome that share a similar profile with the LS elements (Fig. 3D, elements highlighted in black boxes), but as an ensemble, the core elements are statistically different than those found at LS regions.

**Figure 3.**
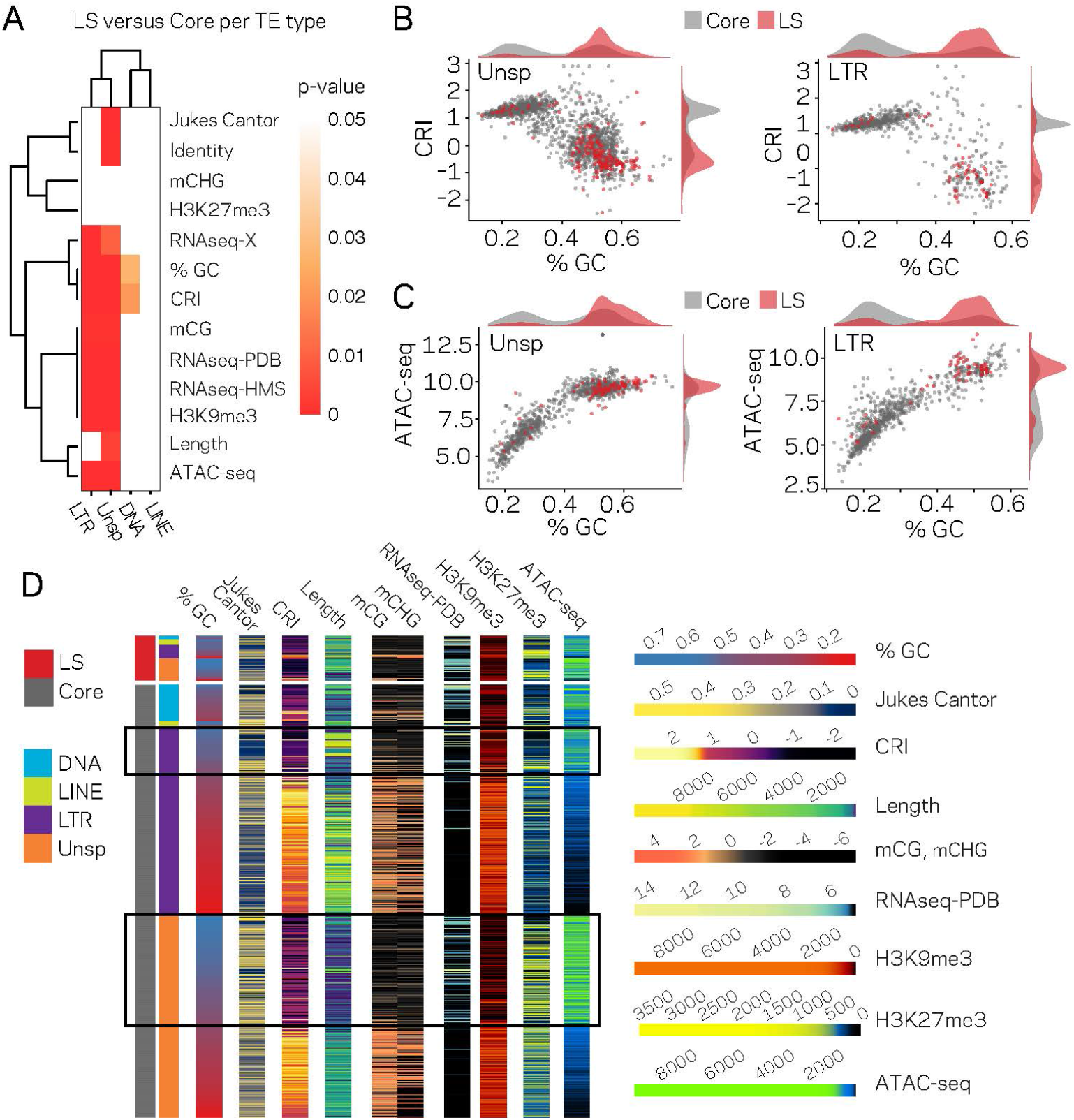
The LTR and Unspecified elements have significantly different chromatin profiles based on core versus LS location. (A) Heatmap comparing core versus LS values within the four TE classifications for the variable listed to the right. Plot colored based on p-values from Wilcoxon rank sum test. P-values ≥ 0.05 are colored white going to red for p-value ≅ 0. (B) Scatter and density plots similar to those shown in Figure 2c except the individual TE points are colored by core (grey) versus LS (red) location. The density plots are also constructed based on the two groupings (C) Similar to B, with the y-axis now showing the log2 (TPM+1) values from ATAC-seq (D) Multiple grouped heatmaps for ten variables collected for each TE. Each row represents a single element and the same ordering is used across all plots. The LS elements are grouped at the top, indicated by the red bar at the top left, and the core elements are grouped below, indicated by the grey bar at the left. Elements are further grouped by the four classifications indicated by the color code shown to the left. Within each element group, the elements are ordered by descending GC content. The scale for each heatmap is shown at the right. % GC, percent GC sequence content; Jukes Cantor, corrected distance as proxy of TE age; CRI, Composite RIP index; Length, element length; mCG and mCHG, log2(weighted cytosine DNA methylation+0.01) for CG and CHG respectively; RNAseq-PDB, variance stabilizing transformed log2 RNA-sequencing reads from PDB grown fungus; H3K9me3 and H3K27me3 and ATAC-seq, TPM values of mapped reads H3K9me3 ChIP-seq, H3K27me3 ChIP-seq, or Assay for transposase accessible chromatin respectively. Black boxes highlight LTR and Unsp elements in the core that have euchromatin profiles.

### Significantly different chromatin status between core and LS regions extends to larger DNA segments

The analysis of TEs in the genome clearly shows that a subpopulation of elements that occur in the previously defined LS regions have different epigenetic modifications and physical openness compared to those in the core genome. LS regions are significant for *V. dahliae* biology as they code many proteins which support host infection. To capture a more global view of core versus LS regions, the genome was analyzed using 10 kb non-overlapping windows, revealing the same global patterns along the linear chromosome sequence; regions with high TE density tend to have lower %GC content, higher DNA and H3K9 methylation and a lack of ATAC-seq reads. The distribution of H3K27me3 appears more complicated. This mark overlaps with that of DNA and H3K9 methylation, as nearly all regions with these two modifications also have H3K27me3, yet we observed additional regions that contain only H3K27me3 and lack DNA and H3K9 methylation (Fig. 4A). The regions that contain DNA methylation and H3K9me3 are nearly identical and for simplicity refer to these regions going forward as being marked by H3K9me3. Interestingly, regions marked by H3K27me3 that lack H3K9me3 have more open DNA than region with H3K27me3 also containing H3K9me3 (Fig. 4A, ATAC). This is apparent for the LS regions that appear to have increased H3K27me3 signal, lack H3K9me3 and are less open than the genomic background but not as closed as the regions marked by H3K9me3 (Fig. 4B, regions marked by grey boxes). PCA was again employed to combine the variables into a single analysis, with the first dimension explaining nearly 60% of the variation in the data (Fig. 4C). The first dimension largely captures the variables describing euchromatin versus heterochromatin, such that ATAC-seq and %GC are furthest separated on the x-axis from H3K9me3, DNA methylation and TE density (Fig. 4C). Interestingly, the DNA segments classified as core are mostly associated with this separation across the first-dimension (Fig. 4C). The second and third dimensions of the PCA explained a similar amount of variation in the data; 14.4% and 10.7%, respectively. Data from the RNA-seq experiment contributed nearly all the information to the second dimension (Supplemental Fig. S4), while the H3K27me3 ChIP-seq data contributed most of the information in the third dimension (Supplemental Table S3). Interestingly, when this third dimension is considered, we observe a strong separation of the core from the LS regions (Fig. 4C, y-axis), suggesting that the LS regions of the genome are less defined by DNA openness, and DNA or H3K9 methylation but more by H3K27me3 and transcriptional activity.

**Figure 4.**
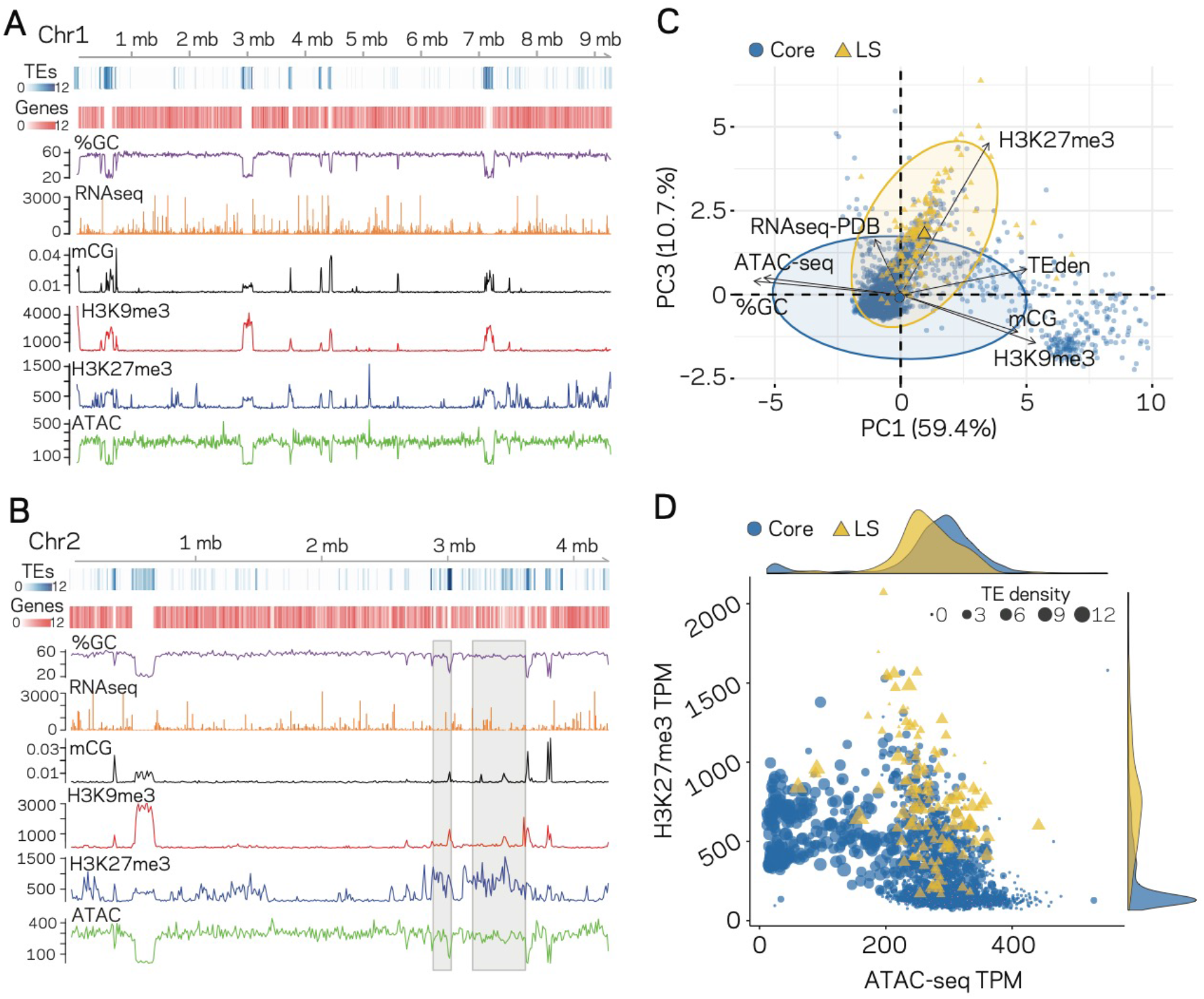
Epigenome and physical DNA characteristics collectively define core and LS regions. (A and B) Whole chromosomes plots showing TE and gene counts over 10 kb genomic windows, blue and red heatmaps respectively. The %GC content is shown in purple, RNA-seq show in orange, CG cytosine DNA methylation shown in black, H3K9me3 and H3K27me3 ChIP-seq shown in red and blue respectively, and ATAC-seq shown in green. Values are those previously described. (B) Two LS regions are highlighted with a grey window. (C) Principle component analysis for seven variables at each 10 kb window. Dimension 1 and 3 are plotted and collective explain ~70% of the variation in the data. The individual symbols are colored by genomic location with core (blue circles) and LS (yellow triangles). Colored ellipses show the confidence interval for the core and LS elements with a single large symbol to show the mean. (D) Scatter plot of the 10 kb windows colored for core and LS location by ATAC-seq data (TPM, x-axis) and H3K27me3 (TPM, y-axis). The size of each symbol is proportional to the TE density shown in the upper right corner. The density plot of each variable is shown on the opposite axis.

Our observations can be summarized into a genome-wide model; for the core genome, regions with higher TE density have low ATAC-seq signal (closed DNA) and elevated H3K9me3 signal and thus represent the heterochromatic regions (Fig. 4D, cluster of large blue dots plotted at middle left). Core genomic regions that are gene-rich have a higher ATAC-seq and lower H3K9me3 signal, and represent the euchromatic portion of the genome (Fig. 4D, cluster of small blue dots plotted in the lower-middle section). The LS regions are a hybrid of the two that contain high TE density and higher H3K27me3 signal but have higher ATAC-seq signals when compared with similar TE containing regions in the core genome (Fig. 4D, cluster of large yellow triangles plotted in the middle). This simple model of the genome accounts for many of the phenomena described here, and links the epigenome, physical genome and functional genome.

### Machine learning predicts more lineage-specific genomic regions than previously considered

Given that a clear model emerges that links the epigenome and physical openness of DNA with adaptive regions of the genome, we assessed the extent to which these features can predict core or LS regions. Stimulated by our observations (Fig. 4), we used ATAC-seq, RNA-seq, H3K27me3, TE density, and H3K9me3 along with the binary classification of the 10 kb windows as core or LS for machine learning. Four supervised machine learning algorithms were used to train (i.e. learn) on 80% of the data (2890 regions), while the remaining 20% (721 regions) were used for prediction (i.e. test), using a 10-fold cross validation repeated three times. Assessing the classifier’s performance using area under the receiver operating characteristic (auROC) curve suggested excellent results ranging from 0.94 to 0.95, with a value of 1 being perfect prediction (Fig. 5A). While auROC is the *de facto* standard for machine learning performance ^59^, it is not appropriate for assessing predictive performance of binary classification problems when the two classes are heavily skewed as it overestimates performance due to the high number of true negatives ^60^. This is the case for our analysis in which the test set (721 regions) contains only 33 of the known LS regions (4.6%). To more accurately assess model performance, precision-recall curves were employed as these do not use true negatives, and are therefore less influenced by skewed binary classifications ^61^. All four algorithms consistently outperformed a random classifier, with the boosted classification tree (BCT) and stochastic gradient boosting (GMB) algorithms having the same highest area under the precision-recall curve of 0.52 (Fig. 5B). However, the confusion matrix indicated that the BCT model only identified 13 of the 33 LS regions (Table 2), resulting in poor recall (Table 3). In contrast, the other three models did identify most of the known LS regions (high recall), but had lower precision caused by the high rate of false positives (Table 2 and 3). The Matthews correlation coefficient (MCC), an analogous measure to accuracy but more appropriate for unbalanced binary classification, indicated that the GMB and random forest (RF) models performed the best (Table 3).

**Figure 5.**
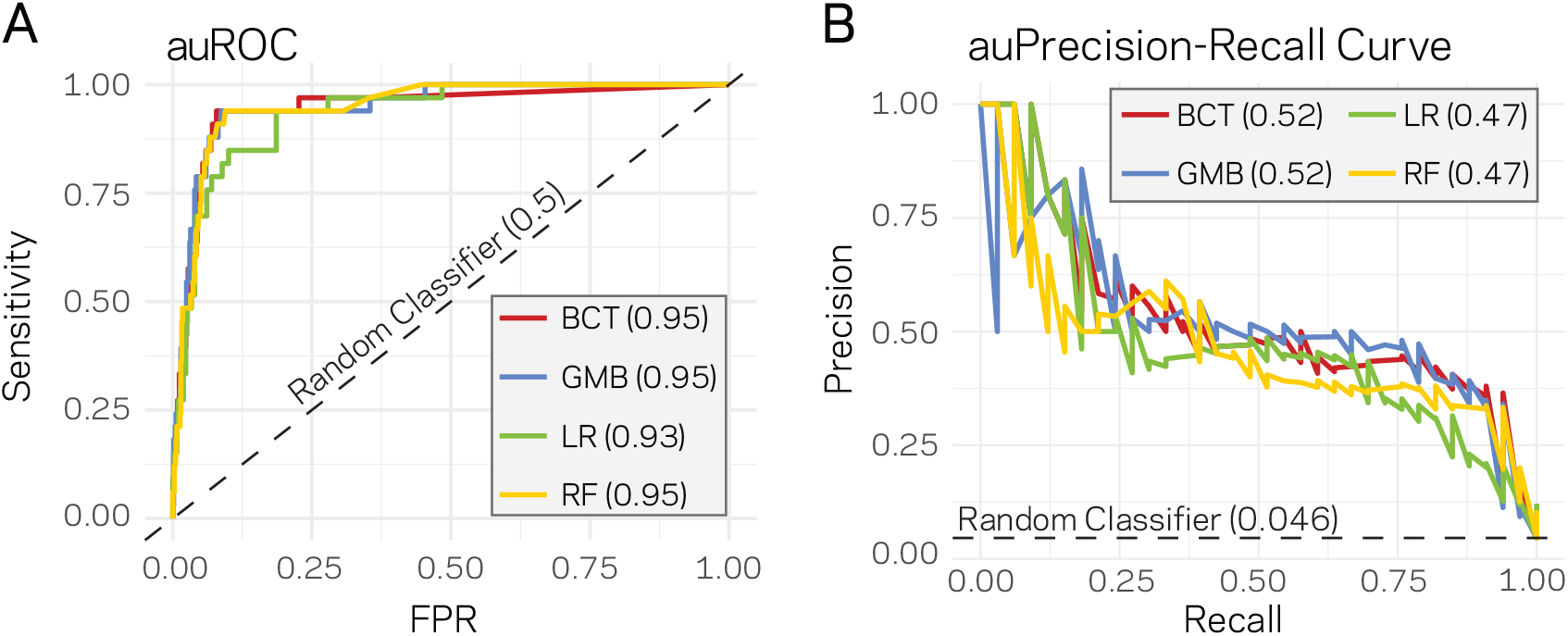
Supervised machine learning can predict LS regions based on epigenome and physical genome characteristics. (A) Area under the Response operator curve (auROC) plotting sensitivity and false positive rate (FPR) for four machine learning algorithms, BCT-Boosted classification tree; GMB-stochastic gradient boosting; LR-logistic regression; RF-random forest. The auROC scores are shown next the algorithm key in the grey box. The black dotted line represents the performance of a random classifier. Perfect model performance would be a curve through point (0,1) in the upper left corner. (B) Area under the Precision-Recall curve for the same four models shown in A. Area under the curves are shown in the figure key in the grey box. The black dashed line shows the performance of a random classifier, calculated as the TP / (TP + FN). Perfect model performance would be a curve through point (1,1) in the upper right corner.

**Table 2.** Confusion Matrix for LS versus core prediction in *V. dahliae*

|  | Predicted | Known |  |
| --- | --- | --- | --- |
|  |  | Core | LS |
| LR | Core | 638 | 7 |
|  | LS | 50 | 26 |
| GMB | Core | 645 | 5 |
|  | LS | 43 | 28 |
| BCT | Core | 672 | 20 |
|  | LS | 16 | 13 |
| RF | Core | 623 | 2 |
|  | LS | 65 | 31 |

LR, Logistic Regression; GMB, Stochastic Gradient Boosting; BCT, Boosted Classification Tree; RF, Random Forest; Core, regions of the genome defined as core; LS, regions of the genome defined as Lineage Specific.

**Table 3.** Assessment values for the four tested machine learning algorithms used to classify genomic regions.

| Models | Precision | Recall | MCC | F1 | F2 |
| --- | --- | --- | --- | --- | --- |
| LR | 0.34 | 0.79 | 0.49 | 0.48 | 0.63 |
| GMB | 0.39 | 0.85 | 0.55 | 0.54 | 0.69 |
| BCT | 0.45 | 0.39 | 0.39 | 0.42 | 0.40 |
| RF | 0.32 | 0.94 | 0.52 | 0.48 | 0.68 |

LR, Logistic Regression; GMB, Stochastic Gradient Boosting; BCT, Boosted Classification Tree; RF, Random Forest; MCC, Matthews Correlation Coefficient.

The results indicate that the machine learning algorithms are well-suited to identify the previously known LS regions in the test data at a high rate. Additionally, the algorithms identified a relatively large number of regions as LS that were previously classified core. The original classification of core and LS in *V. dahliae* was based on presence/absence variations identified from genomic information of only few strains ^14,15^. Consequently, we reasoned that regions here classified as LS by the machine learning algorithms could be genuine LS regions that were originally missed due to the limited diversity of the *V. dahliae* represented by the strains sequenced. The two best models from the initial testing, GMB and RF, predicted a total of 96 and 81 regions as LS respectively, suggesting there could be 2 to 3 times more LS DNA than previously identified. To improve the genome-wide estimate and to further assess the robustness of machine learning for LS region prediction, we re-ran the GMB and RF algorithms on 15 new training-test splits, independently training and predicting on each set (see methods for details). This approach nearly saturated the genome, providing multiple predictions per window and only 124 of the 3611 regions were missed (Supplemental Fig. S5). The average MCC performance estimate of the GMB and RF classifiers were 0.53 and 0.48 over the 15 runs, and our results indicate consistent performance across the independent predictions (Fig. 6A, Supplemental Fig. S6, Supplemental Table S4 and S5). The GMB classifier predicted a total of 285 of the 10 kb regions as LS, while the RF classifier predicted 388 (Supplemental Table S6 and S7). The LS predictions for the two models were in agreement for 280 regions, which is 98% of the GMB predictions and 72% of those from the RF (Fig. 6B), overall showing high agreement between the two classifiers. Consensus predictions were generated from the two classifiers if a region was predicted as LS by both models, and a conservative joining step was employed in which a single predicted core region was called LS if it was flanked by LS predictions on both sides (see methods). This resulted in a total of 280 regions predicted as LS by both classifiers and an additional 41 regions due to the joining. In total, this new classification nearly doubles the total amount of LS regions compared with the original observations ^14,15^. The original classification of LS regions in *V. dahliae* clustered in four larger regions^14,15^. We were interested to understand the physical genomic location of the originally identified and the newly predicted LS regions. The results of the individual classifiers reveal that the new regions are also not randomly dispersed across the genome (Supplemental Fig. S7). The consensus prediction from the two classifiers identified the large blocks of LS regions from the original observations, along with new clusters of LS regions such as those on chromosomes 4, 6, and 8 (Fig. 6C and 6D). Importantly, the newly defined set of LS regions supports a clearer separation of the LS regions from the core regions (Supplemental Fig. S8). Collectively, these analyses suggest that the machine learning algorithms can be used to predict new LS regions based on epigenetic and physical DNA accessibility data. The identification of potentially new LS regions missed in the original classification provides new avenues to identify proteins important for host infection and adaptation. These results support that genome structure is influencing genome function, demonstrates a machine learning approach for predictive biology, and advances our biological understanding of genome function.

**Figure 6.**
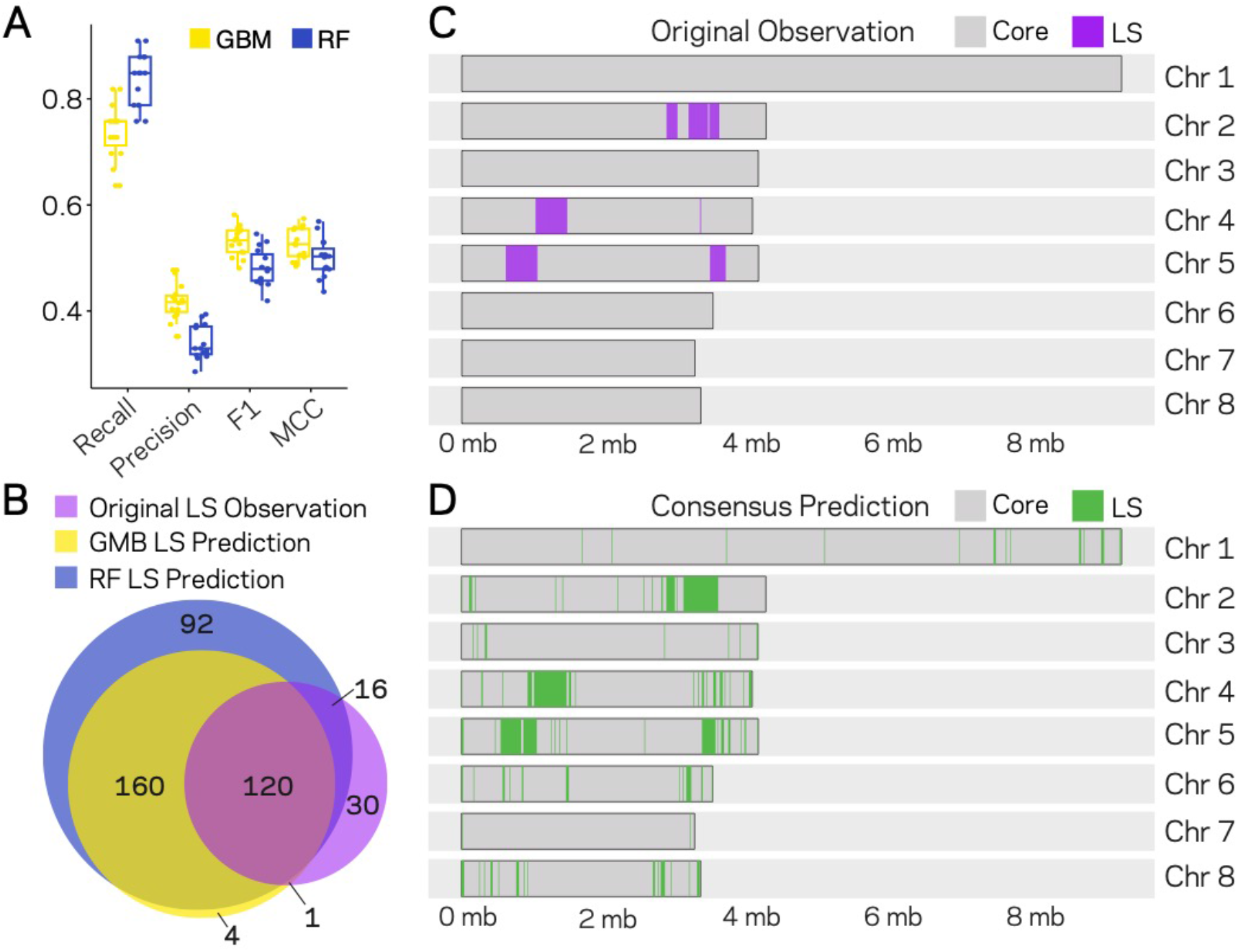
Machine Learning predictions for genome-wide LS content. (A) Two machine learning algorithms, Stochastic Gradient Boosting (GMB) and Random Forest (RF), were used to predict LS regions from 15 independent training-test splits. For each split, 80% of the data were used to train and the remaining 20% were used for prediction. Classifier performance was measured for each of the 15 trials, and summarized as a boxplot with each trial represented as a point. (B) Venn diagram showing the overlap between the results of the two classifiers and the original observations of LS regions (Faino et al. 2016). Each slice of the diagram shows the number of LS regions predicted, see methods for additional details. (C and D) Schematic representation of the eight chromosomes (labeled on right) of *V. dahliae* strain JR2. Each chromosome was divided into 10 kb windows for prediction. Regions classified as core are shown in grey and LS as their indicated color. (C) Original observations of core and LS regions, grey and purple respectively. The five main LS regions 1can be seen on chromosome 2, 4 and 5. (D) The consensus model predictions for core and LS regions shown in grey and green respectively. The consensus predictions were those made by both the GMB and RF model (in total 280). Regions predicted as LS were joined if they were interrupted by a single window of Core prediction, adding an additional 41 LS regions for a final number of 321 LS regions in the *V. dahliae* JR2 genome.

## DISCUSSION

Significant efforts to detail genomes of filamentous pathogens, to understand variation within species, and to a lesser extent to examine epigenetic modifications, have increased our understanding genome function in this important group of organisms ^16,54,62^. Understanding pathogen genome evolution is of great interest to help combat emerging pathogens, and to broaden our knowledge of genome biology beyond model eukaryotes. Here we present a detailed analysis of the epigenome and physical DNA accessibility of the vascular wilt pathogen *V. dahliae* and link these analyses to previous characterizations of genomic regions contributing to host colonization and adaptation ^13–16^. A clear picture emerges in which the core genome is organized into heterochromatic and euchromatic regions. The heterochromatin is characterized by a high density of TEs with low GC content, high levels of DNA and H3K9 methylation, low DNA accessibility and clear signatures of RIP mutations at repetitive sequences. The euchromatin regions are opposite in all characteristics, and this collective description is consistent with previous research in many other eukaryotic genomes ^32,63,64^. Interestingly, we provide evidence that previously defined LS regions of the genome, characterized for their role in contributing to host infection, exist in an intermediate chromatin state, having higher TE density than the euchromatic regions, yet are devoid of DNA and H3K9 methylation. Furthermore, LS regions have higher DNA accessibility than the core heterochromatic regions and are more transcriptionally active, but they are less accessible than the ‘true’ euchromatic gene-rich core regions. Notably, LS regions are characterized as having a strong association with H3K27me3, similar to the discovery that SM gene clusters are enriched at H3K27me3 regions in *F. graminearum* ^29^. Our results demonstrate that LS regions are by definition not heterochromatic, as they are far more accessible than the true heterochromatin, and yet they typically contain many heterochromatin features. We note previous descriptions of contradictory heterochromatin states ^65^, and the broad possible chromatin states that may characterize a genome ^33^. However, few previous analyses have assessed the relationship between DNA and histone modifications with DNA accessibility in light of biological function of genomic adaptation.

Our results support the hypothesis that chromatin structure underlies genome function. More specifically that chromatin modifications and DNA accessibility contribute to genome evolution, not just via transcriptional control but also regarding the architecture of the genome ^50^. Along with the described associations, we were able to predict LS regions using machine learning. The results of running four machine learning algorithms trained on H3K9 and H3K27 methylation, RNA-sequencing, TE density and DNA accessibility data, shows these variables could be used to classify DNA segments as core versus LS with high recall (i.e. sensitivity). The RF model showed the highest recall, correctly classifying 31 of the previously observed 33 LS regions in spite of their skewed presence in the data at nearly 1:20 LS to core. The precision assessment of the algorithms was low because each model classified regions as LS that were originally observed as core, statistically termed false positives. However, the original observations represent operational classification based on then available data. Consensus predictions based on the two highest performing models extended the boundaries of the previous LS regions and identified new potentially clustered LS regions. Thus, the use of machine learning can extend our knowledge of biology and identify novel genomic regions to search for as of yet uncharacterized genes with important adaptive roles. Collectively, we interpret our results to indicate a strong link between the epigenome, physical DNA accessibility and the occurrence of LS regions in *V. dahliae*. Our findings however have limited inference on causation versus association, an important area for future research. If there is a causative relationship between genome structure and function it is interesting to consider who drives whom- do the LS regions dictate altered chromatin or does altered chromatin dictate LS formation?

It is currently not possible to extend our machine learning predictions to additional filamentous pathogen genomes, as the necessary data are not currently publicly available. However, for many filamentous plant pathogens it is clear that genome variation on multiple scales, from SNPs to large structural variation, are not uniformly distributed in the genome ^29^. Recent reports from the fungal pathogen *Z. tritici* addressed the role of genome stability and H3K27me3 during asexual reproduction ^53,66^. During experimental evolution, individual strains of *Z. tritici* readily lose accessory chromosomes. The authors observed that a mutant lacking the enzyme responsible for H3K27me3 showed less accessory chromosome loss and concluded that H3K27me3 destabilizes chromosome structure ^53^. However, accessory chromosome losses were clearly biased in their individual frequency and changes were not reported for core chromosomes, despite H3K27me3 being found at high levels on accessory and regions of core chromosomes ^67^. Therefore, the observed genome destabilization requires additional determinants in conjunction with H3K27me3 which remain to be discovered. Results presented here suggest that DNA and histone methylation marks and physical DNA accessibility are important additional determinants to distinguish accessory and LS regions of the genome. However, we acknowledge that our model does not strictly differentiate all LS region in the *V. dahliae* genome, as there are LS and core regions that have very similar overall chromatin profiles, and therefore these features alone are not sufficient. One factor that could explain part of this discrepancy is that LS formation is likely not fully deterministic. Evolution is a stochastic process, and it seems unlikely that LS formation can be described in absolute terms. Rather, it is more likely to be a probabilistic process, in which specific chromatin and physical status increases the likelihood for formation and maintenance of LS regions. The results presented here offer an exciting new link between the epigenome, physical DNA accessibility and adaptive genome evolution.

## METHODS

### Fungal growth and strain construction

*V. dahliae* strain JR2 (CBS 143773) was used for experimental analysis ^68^. The strain was maintained on potato dextrose agar (PDA) (Oxoid, Thermo Scientific, CM0139) and grown at 22⁰C in the dark. For liquid grown cultures, conidiospores were collected from PDA plates after approximately two weeks and inoculated into flasks containing the desired media at a concentration of 2×10^4^ spores per mL. Media used in this study include PDA, half-strength Murashige and Skoog plus vitamins (HMS) (Duchefa-Biochemie, Haarlem, The Netherlands) medium supplemented with 3% sucrose and xylem sap (abbreviated, X) collected from greenhouse grown tomato plants of the cultivar Moneymaker. Liquid cultures were grown for four days in the dark at 22⁰C and 160 RPM. The cultures were strained through miracloth (22 μm) (EMD Millipore, Darmstadt, Germany), pressed to remove liquid, flash frozen in liquid nitrogen and ground to powder with a mortar and pestle. Samples were stored at −80⁰C if required prior to nucleic acid extraction.

The *Δhp1* strain was constructed as previously described ^69^. Briefly, the genomic DNA regions flanking the 5’ and 3’ HP1 coding sequence were amplified (*left border*, For. Primer, 5’-GGTCTTAAUGACCTGAAGAATCGAGCAAGGA and Rev. primer, 5’-GGCATTAAUATGAAAGCACCGGGATTTTTCT; *right border*, For. Primer, 5’-GGACTTAAUATGCTGTTGGGAGGCAGAATAA Rev. primer, 5’-GGGTTTAAUCCACGTAGATGGAGGGGTAGA). The PCR products were cloned in to the pRF-HU2 vector system ^70^ using USER enzyme following manufactured protocol (New England Biolabs, MA, USA). Correctly ligated vector was transformed into *Agrobacterium tumefaciens* strain AGL1 used for *V. dahliae* spore transformation ^69^. Colonies of *V. dahliae* growing on hygromycin B selection after 5 days were moved to individual plates containing PDA and hygromycin B. Putative transformants were screened using PCR to check for deletion of the HP1 sequence (For. Primer, 5’- AATCCCGCAAGGGAAAAGAGAC and Rev. primer, 5’- CGTGTGCTTTGTCTTCTGACCA) and the integration of the hygromycin B sequence (For. Primer, 5’- TGGAATATGCCACCAGCAGTAG and Rev. primer, 5’- GGAGTCGCATAAGGGAGAGCG) at the specific locus.

### Bisulfite sequencing and analysis

The wild-type *V. dahliae* strain and *Δhp1* were grown in liquid PDA for three days, flash frozen and collected as described earlier. Extracted DNA was sent to the Beijing Genome Institute (BGI) for bisulfite conversion, library construction and Illumina sequencing. Briefly, the DNA was sonicated to a fragment range of 100-300 bp, end-repaired and methylated sequencing adapters were ligated to 3’ ends. The EZ DNA Methylation-Gold kit (Zymo Research, CA, USA) was followed according to manufacturer guidelines for bisulfite conversion of non-methylated DNA. Libraries were paired-end 100bp sequenced on an Illumina HiSeq 2000.

Whole-genome bisulfite sequencing reads were analyzed using the BSMAP pipeline (v. 2.73) and methratio script ^71^. The results were partitioned into CG, CHG and CHH cytosine sites for analysis. Only cytosine positions containing greater than 4 sequencing reads were included for analysis. Methylation levels were summarized as weighted methylation percentage, calculated as the number of reads supporting methylation over the number of cytosines sequenced or as fractional methylation, calculated as the number of methylated cytosines divided by all cytosine positions^72^. For fractional methylation, a cytosine was considered methylated if it was at least 5% methylated from all the reads covering that cytosine. As such, weighted methylation captures quantitative aspects of methylation, while fractional methylation is more qualitative. Weighted and fractional methylation were calculated over intervals described in the text, including genes, promoters (defined as the 300 bp upstream of the translation start site), transposable elements and 10 kb windows. For each feature, weighted and fractional methylation were calculated from the sum of the mapped reads or the sum of the positions, respectively, over the analyzed region. Two sample comparisons were computed using base R ^73^ to compute the non-parametric Mann-Whitney U test (equivalent to the two-sample Wilcoxon rank-sum test). Principle component analyses were computed in R using the packages FactoMineR (v 1.42) ^74^ and factoextra (v 1.0.5) ^75^.

### Transposable element annotation

Repetitive elements were identified in the *V. dahliae* stains JR2 genome assembly ^68^ as well as in three other high-quality *V. dahliae* genome assemblies ^16^ using a combination of LTRharvest ^76^ and LTRdigest ^77^ followed by *de novo* identification of RepeatModeler ^78^. Briefly, LTR sequences were identified (recent and ancient LTR insertions) and subsequently filtered, e.g. for occurrence of primer binding sites or for nested insertions (see procedure outlined by Campbell and colleagues for details^79^). Prior to the *de novo* prediction with RepearModeler, genome-wide occurrences of the identified LTR elements are masked. Predicted LTR elements and the *de novo* predictions from RepeatModeler were subsequently combined, and the identified repeat sequences of the four *V. dahliae* strains were clustered using vsearch (80% sequence identity, search on both strands; v 2.4.4) ^80^. A non-redundant *V. dahliae* repeat library that contained consensus sequences for each cluster (i.e. repeat family) was constructed by performing multiple sequence alignments using MAFFT (v7.271) ^81^ followed by the construction of a consensus sequence as described by Faino et al. ^15^. The consensus repeat library was subsequently manually curated and annotated (Wicker classification ^82^) using PASTEC (default databases and settings; search in the reverse-complement sequence enabled) ^83^, which is part of the REPET pipeline (v2.2) ^84^, and similarity to previously identified repetitive elements in *V. dahliae* ^68,85^. The occurrence and location of repeats in the genome assembly of *V. dahliae* strain JR2 were determined using RepeatMasker (v 4.0.7; sensitive option). The Repeatmasker output was post-processed using ‘One code to find then all’ ^86^ which supports the identification and combination of multiple matches (for instance due to deletions or insertions) into combined, representative repeat occurrences. We only further considered matches to the repeat consensus library, and thereby excluded simple repeats and low-complexity regions. To estimate divergence time of TEs, each individual copy of a transposable element was aligned to the consensus of its family using needle, which is part of the EMBOSS package ^87^. Sequence divergence between the TEs and the TE-family consensus was corrected using the Jukes-Cantor distance, with a correction term that accounts for insertions and deletions ^88,89^. The composite RIP index (CRI) was calculated as previously described ^43^. Briefly, CRI was determined by subtracting the RIP substrate from the RIP product index, which are defined by dinucleotide frequencies as follows: RIP product index = (TpA / ApT) and the RIP substrate index = (CpA + TpG/ ApC + GpT). Positive CRI values indicate the analyzed sequences were subjected to the RIP process. For TE analysis, elements that are less than 100 bp were removed.

### RNA-sequencing and analysis

*V. dahliae* strain JR2 (CBS 143773) was grown in triplicate liquid media PDB, HMS and xylem sap as described. RNA extraction was carried out using TRIzol (Thermo Fisher Science, Waltham, MA, USA) following manufacturer guidelines. Following RNA re-suspension, contaminating DNA was removed using the TURBO DNA-free kit (Ambion, Thermo Fisher Science, Waltham, MA, USA) and RNA integrity was estimated by separating 2 μL of each sample on a 2% agarose gel and quantified using a Nanodrop (Thermo Fisher Science, Waltham, MA, USA) and stored at −80⁰C. Library preparation and sequencing was carried out at BGI. Briefly, mRNA were enriched based on polyadenylation purification and random hexamers were used for cDNA synthesis. RNA-sequencing libraries were constructed following end-repair and adapter ligation protocols and PCR amplified. Purified DNA fragments were single-end 50bp sequenced on an Illumina HiSeq 2000.

Reads were mapped to the *V. dahliae* stain JR2 genome assembly ^68^ using STAR (v 2.6.0) with settings *(--sjdbGTFfeatureExon exon, --sjdbGTFtagExonParentTranscript Parent, --alignIntronMax 400, --outFilterMismatchNmax 5, --outFilterIntronMotifs RemoveNoncanonical*) ^90^. Mapped reads were quantified using the *summarizeOverlaps* and variance stabilizing transformation (*vst*) features of DESeq2 ^91^. For TE analysis, the coordinates of the annotated TEs were used as features for read counting. To perform RNAseq analysis over whole genome 10 kb regions, raw mapped reads were summed over 10 kb bins using bedtools (v 2.27) ^91^ and converted to Transcripts Per Million (TPM) and averaged over the three reps for analysis.

### Chromatin immunoprecipitation and sequencing and analysis

*V. dahliae* strain JR2 was grown in PDB and materials was collected as described. Approximately 400 mg ground material was resuspended in 4 ml ChIP Lysis buffer (50 mM HEPES-KOH pH7.5, 140 mM NaCl, 1 mM EDTA, 1% Triton X-100, 0.1% NaDOC) and dounced 40 times in a 10 cm^3^ glass tube with tight fitting pestle on 800 power with a RZR50 homogenizer (Heidolph, Schwabach, Germany), followed by five rounds of 20 seconds sonication on ice with 40 seconds rest between rounds with a Soniprep 150 (MSE, London, UK). Samples were redistributed to 2 ml tubes and pelleted for 2 min at max speed in tabletop centrifuge. The supernatants were combined, together with 20 µl 1M CaCl2 and 2.5µl MNase, and after 10 minutes of incubation in a 37°C water bath with regular manual shaking, 80 µl 0.5M EGTA was added and tubes were put on ice. Samples were pre-cleared by adding 40 µl Protein A Magnetic Beads (New England Biolabs, MA, United States) and rotating at 4°C for 60 min, after which the beads were captured, 1 ml fractions of supernatant were moved to new 2 ml tubes containing 5 μl H3K9me3 or H3K27me3 antibody (ActiveMotif; #39765 and #39155) respectively and incubated overnight with continuous rotation at 4ᵒC. Subsequently, 20 μl protein-A magnetic beads were added and incubated for 3 hours at 4ᵒC, after which the beads were captured on a magnetic stand and subsequently washed with 1 ml wash buffer (50 mM Tris HCl pH 8, 1 mM EDTA, 1% Triton X-100, 100 mM NaCL), high-salt wash buffer (50 mM Tris HCl pH 8, 1 mM EDTA, 1% Triton X-100, 350 mM NaCL), LiCl wash buffer (10 mM Tris HCl pH8, 1 mM EDTA, 0.5% Triton X-100, 250 mM LiCl), TE buffer (10 mM Tris HCl pH 8, 1mM EDTA). Nucleosomes were eluted twice from beads by addition of 100μl pre-heated TES buffer (100 mM Tris HCl pH 8, 1% SDS, 10 mM EDTA, 50 mM NaCl) and 10 minutes incubation at 65ᵒC. 10mg /ml 2μl Proteinase K (10mg /ml) was added and incubated at 65ᵒC for 3 hours, followed by chloroform clean-up. DNA was precipitated by addition of 2 volumes 100% ethanol, 1/10th volume 3 M NaOAc pH 5.2 and 1/200th volume 120 mg/ml glycogen, and incubated overnight at −20ᵒC. Sequencing libraries were prepared using the TruSeq ChIP Library Preparation Kit (Illumina) according to instructions, but without gel purification and with use of the Velocity DNA Polymerase (BioLine, Luckenwalde, Germany) for 25 cycles of amplification. Single-end 125bp sequencing was performed on the Illumina HiSeq2500 platform at KeyGene N.V. (Wageningen, the Netherlands).

Reads were mapped to the reference JR2 genome, using BWA-mem with default settings ^92^. For ChIP and ATAC-seq mapping, three regions of the genome were masked due to aberrant mapping, possibly owing to sequence similarity to the mitochondrial genome (chr1:1-45000, chr2:3466000-3475000, chr3:1-4200). This is similar to what is described as blacklisted regions in other eukaryotic genomes ^93^. The raw mapped reads were counted either over the TE coordinates or 10 kb intervals for the two separate analyses. The raw mapped reads were converted to TPM and the average of the two replicates was used for analysis.

### Assay for Transposase-Accessible Chromatin (ATAC)-sequencing and analysis

The *V. dahliae* strain JR2 (CBS 143773) was grown in PDB liquid media as described. Mycelium was collected, filtered, rinsed and flash frozen in liquid nitrogen. The ATAC-seq procedure was carried out mainly as described previously ^94^. Nuclei were collected by resuspending ground mycelium in 5 mL of ice-cold Nuclei Isolation Buffer (NIB) (100 mM NaCl, 4mM NaHSO_4_, 25mM Tris-HCl, 10mM MgSO_4_, 0.5mM EDTA, 0.5% NP-40 including protease inhibitors added at time of extraction, 2 mM Phenylmethanesulfonyl fluoride (PMSF), 100 µM Leupeptin, 1 µg/mL Pepstatin, 10 µM E-64). The homogenate was layered onto 10-mL of an ice-cold sucrose-Ficoll gradient (bottom layer 5mL of 2.5M sucrose in 25mM Tris-HCl, 5mL 40% Ficoll 400 (GE Biosciences Corporation, NJ, USA)). Nuclei were separated into the lower phase by centrifugation at 2000g for 30 min at 4ᵒC. The upper layer was discarded and the lower phase (~4mL) moved to another collection tube containing 5mL of ice-cold NIB. Nuclei were pelleted at 9000g for 15 min at 4ᵒC and re-suspended in 3 mL of NIB. The integrity of the nuclei and their concentration in the solution were estimated by DAPI staining (DAPI Dilactate 5mg/mL, used at a 1/2000 dilution for visualization) and counted on a hemocytometer. A total of 200,000 nuclei were transferred to a 1.5mL microfuge tube, and nuclei pelleted at 13000g for 15 min at 4ᵒC and resuspended in the transposition reaction (20uL of 2x Nextera reaction buffer, 0.5uL of Nextera Tn5 Transposase, 19.5 uL of nuclease-free H_2_0) (Illumina, Nextera DNA library Preparation kit FA-121-1030) and the reaction was carried out for 5 minutes at 37ᵒC. The reaction was halted and fragmented DNA purified using a MinuElute PCR purification kit (Qiagen, MD, USA). The eluted DNA was amplified in reaction buffer (10uL of transposased DNA, 10uL nuclease-free H_2_0, 2.5uL forward PCR primer (5’-AATGATACGGCGACCACCGAGATCTACACTCGTCGGCAGCGTCAGATGTG), 2.5uL reverse PCR primer (CAAGCAGAAGACGGCATACGAGATTTCTGCCTGTCTCGTGGGCTCGGAGATGT) and 25uL NEBnext High-Fidelity 2x PCR Master Mix (New England Biolabs, MA, United States)) using thermo-cycler conditions described in ^94^ for a total of 9 cycles. Amplified library was purified using the MinElute PCR Purification Kit (Qiagen, MD, USA) and paired-end 100 bp sequenced on an Illumina HiSeq4000.

Reads were mapped to the reference JR2 genome with the described blacklisted regions masked, using BWA-mem with default settings ^92^. The mapped reads were further processed to remove duplicates reads arising from library prep and sequencing using Picard toolkit *markDuplicates* ^95^. The mapped reads were counted either over the TE coordinates or 10 kb intervals for the two separate analyses using bedtools *multicov* (v 2.27) ^96^. The reads were converted to TPM values and those numbers used for analysis.

### Machine Learning and assessment

The machine learning algorithms were implemented using the classification and regression training (caret) package in R ^73,97^. The full set of genomic data was used to create a data frame comprising the genome in 10 kb segments as rows and the individual collected variables as columns. The regions were classified as core or LS based on the previous observations ^15^. For initial model assessment and parameter tuning, the data were split into 80% for training and 20% used for testing (i.e. prediction), and the proportion of core and LS regions were kept approximately equal in the two splits. For parameter tuning, repeated cross-validation of 10-fold 3- times was used and the best model was selected based on accuracy. Four algorithms were used-logistic regression, random forest, stochastic gradient boosting, and boosted classification tree. The model for all algorithms was classification = ATAC-seq_TPM_ + ChIP-H3K27me3_TPM_ + ChIP-H3K9me3_TPM_ + TE_density_ + PDB-RNAseq_TPM_. Logistic regression was run using method *glm*, family *binomial*. Random forest was run using method *rf* and tuneGrid [*mtry*= (1,2,3)]. The Stochastic Gradient Boosting was implemented with method *gbm* and tuneGrid [*interaction.depth*=(1,5,10), *n.trees*=(50,500,1000), *shrinkage*=(0.001, 0.01), *n.minobsinnode*=(1,5)]. The Bosted Classification Tree was implemented unsing method *ada* and tuneGrid [*iter*=(100, 1000, 3000), *maxdepth*=(1,5,20), *nu*=(0.01)]. Models were assessed using standard metrics for data retrieval, with receiver operating and precision-recall curves generated using package PRROC ^98^.

To saturate the genome in predictions, a total of 15 new training test splits (80:20) were generated, again maintaining the genome-wide proportion of core and LS regions in data set. The random forest and stochastic gradient boosting classifiers were used, based on their highest performance from the initial test. The settings were picked based on best performance from initial testing: random forest, method *rf* and tuneGrid [*mtry*=3]; stochastic gradient boosting, method *gbm* and tuneGrid [*interaction.depth*=(5), *n.trees*=(500), *shrinkage*=(0.01), *n.minobsinnode*=(5)]. The predictions for each of the 15 runs were assessed using the precision, recall and MCC metrics. For each genomic region, a consensus designation was assigned based on the highest occurrence of core versus LS prediction across the 15 trials. This was done independently between the two models. A region was finally classified as LS or core based on the majority classification across the 15 trails. For regions that had an equal number of core and LS predictions, the regions were designated as core to be conservative. A final high confidence LS consensus designation was determined for each genomic region if it was predicted LS by both models. Regions predicted LS by only one of the models were designated core. A conservative joining approach was used so that a single core region would be called LS if it were flanked by two LS regions. This added 41 genomic regions (410 kb) to the LS genome.

## DATA ACCESS

The sequencing data for this project are accessible from the National Center for Biotechnology Information (NCBI) Sequence Read Archive (SRA) under BioProject PRJN592220.

## ACKNOWLEDGEMENTS

This work was supported in part by a European Molecular Biology Organization postdoctoral fellowship (EMBO, ALTF 969-2013) and Human Frontier Science Program Postdoctoral Fellowship (HFSP, LT000627/2014-L) to DEC. A portion of the work was also carried out in the Cook lab under USDA-NIFA-PBI grant (2018-67013-28492). Work in the laboratories of M.F.S and B.P.H.J.T. is supported by the Research Council Earth and Life Sciences (ALW) of the Netherlands Organization of Scientific Research (NWO).

## SUPPLEMENTAL MATERIAL

Supplemental Fig S1

Supplemental Fig S2

Supplemental Fig S3

Supplemental Fig S4

Supplemental Fig S5

Supplemental Fig S6

Supplemental Fig S7

Supplemental Fig S8

Supplemental Table S1

Supplemental Table S2

Supplemental Table S3

Supplemental Table S4

Supplemental Table S5

Supplemental Table S6

Supplemental Table S7

